# Modeling vaccine-induced immunotherapy: treatment scheduling and robustness with virtual mice cohorts

**DOI:** 10.1101/740878

**Authors:** Priya Bhatt, Manoj Kambara, Shari Pilon-Thomas, Katarzyna A. Rejniak, Ibrahim M. Chamseddine

**Affiliations:** Ottawa Hills High School. Ottawa Hills, OH; Land O’Lakes High School, Land O’Lakes, FL; The Integrated Mathematical Oncology Department; The Immunology Department; H. Lee Moffitt Cancer Center & Research Institute, Tampa FL 33612

## Abstract

Therapeutic vaccines are used to boost patients’ immune system activity by imposing signals that increase T cell proliferation or infiltration. A large population of cytotoxic T cells may then be able to reduce tumor growth. We developed here a mathematical model of vaccine-induced immunotherapy and used it to test the vaccine frequency and doses that can reduce tumor burden. Since tumors are heterogeneous, we examined if the proposed treatments are robust; i.e, are successful for a wide range of tumors. This was assessed by constructing virtual mice cohorts. Together, the optimal and most robust treatment protocol was determined through mathematical modeling.

## I. INTRODUCTION

The patient’s own immune system provides the first line of defense against foreign bodies, such as viruses or cancer cells. However, immune cells are rarely effective in fight with larger tumors. One of the methods to boost patients’ immune system activity is administration of therapeutic cancer vaccines [1]. Vaccines within the tumor tissue transfect tumor cells which leads the immune system to bring in more T-cells to the tumor site, and also increase the proliferation of those T-cells at the tumor site. This boost in the number of T-cells will allow for further destruction of the tumor cells by T-cells. However, tumor cells can suppress T cell activity. Therefore, further studies to optimize vaccination dosage and timing, as well as testing how well different tumors will respond to these treatments are needed.

## II. DESIGNING EFFECTIVE TREATMENT SCHEDULES

Various treatment protocols were simulated to find the potential optimal treatment (Figure 1). A total of 60 mcg of vaccine was administered in our mathematical model in equally spaced injections throughout the treatment period of 25 days. The simulations included protocols with a single 60 mcg injection at day 7; two 30 mcg injections and days 7 and 16; three 20 mcg injections at days 7, 13 and 17; and four 15 mcg injections at days 7, 12,16 and 21 (Figure 1C).

**Figure 1.**
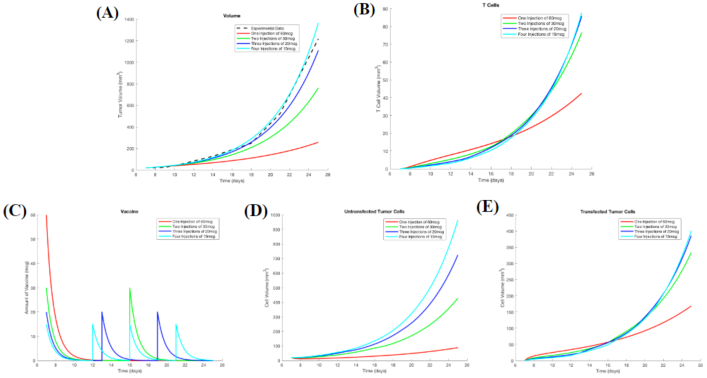
Four treatment schedules with 1 (red), 2 (green), 3 (blue) and 4 (cyan) vaccine injections: (A) The total tumor volume; (B) The volume of T cells on the tumor site; (C) Vaccine volume at the tumor site. (D) Untransfected tumor cell volue; (E) Transfected tumor cell volume.

Figure 1A indicates that the tumor growth is suppressed the most significantly when a single, large injection of 60 mcg of vaccine is given early, at day 7. This is supported by Figure 1B, which indicates that the single injection yielded the greatest amount of T cells at the tumor site at the beginning of treatment, when the tumor volume is still small enough to be controlled. Moreover, the ratio of transfected (Figure 1E) to untransfected (Figure 1D) tumor cells is higher in the case of the single vaccine injection when compared to all other protocols.

Using this mathematical model, we investigated a wide variety of treatment protocols, but we continued to find that a single, large early dose yielded optimal results.

## III. GENERATING VIRTUAL MICE COHORTS

Because no two tumors are the same, treatment schedules should be tested on a broad range of individual data to account for tumor heterogeneity and assess the robustness of the therapy. When a treatment is deemed as robust, it can reach many heterogeneous individuals with the desired effect.

To test the robustness of treatment protocols, we created virtual mice cohorts by adapting the method described in Barish et. al. [2]. Three key model parameters were chosen: tumor cell net proliferation (r), tumor cell transfection rate (β), and tumor cell killing rate by the T cells (k), and the generated distributions for each of these parameters are shown in Figure 2.

**Figure 2.**
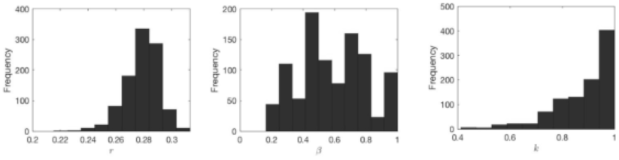
Frequency distributions for (left) tumor proliferation rate *r*, (middle) tumor transfection rate *β*, and (right) tumor killing rate (k), generated using the virtual expansion procedure.

By sampling parameters within these distributions, 1,000 heterogeneous virtual mice cohorts were created. Finally, these cohorts were used to test outcomes of various scheduling procedures and assess their robustness.

## IV. TESTING TREATMENT ROBUSTNESS

Each of the four treatment schedules from Section II were tested against the same 1,000 virtual mice cohorts, and the tumor size after simulated 25 days of treatment was recorded. These measurements were used to determine the mice cohort objective response to a given schedule and schedule ranking for a give cohort.

The objective response of a mouse was marked positive (tumor response), if the mouse yielded a tumor area lower than the predefined expected average (137 mm^2^ in area), negative (non-response) otherwise. Moreover, we marked which treatment yielded the lowest tumor area, essentially ranking which of the four treatments was the best for each mouse. The frequency of these measurements is then collected for each treatment as shown in Figure 3. Each bar represents what portion of the 1,000 virtual mice responded to the treatment.

**Figure 3.**
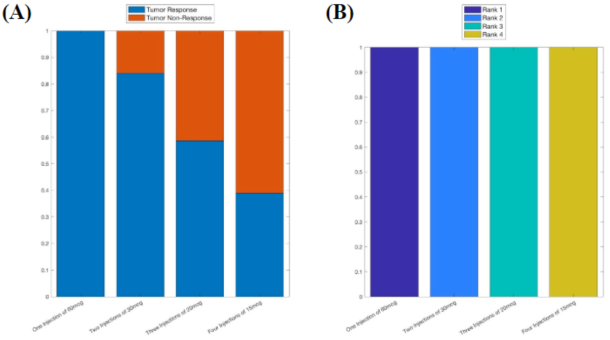
(A) The frequency at which the entire virtual population had an objective response. (B) The frequency at which the treatment ranked for the virtual population. Higher rank implies lower tumor area. The bars represent 4 differnet treatments; from left to right: 1, 2, 3 or 4 injections.

The virtual cohort simulations showed that a single 60 mcg injection seemed to yield the highest frequency of tumor response (Figure 3A), and for each mouse this treatment yielded the lowest tumor area resulting in 100% rank 1 (Figure 3B). Though other treatments also seem to show robust results such as the two-injection schedule, the single injection yielded the most robust response. This means, the single, early injection will yield the best results for a heterogeneous population with tumors.

## V. QUICK GUIDE TO THE EQUATIONS

The mathematical model describes temporal changes in the volumes of four species: untransfected tumor cells (U), transfected tumor cells (I), the vaccine (V), and T cells (T). The change from untransfected to transfected tumor cells is triggered by the vaccine, which upon transfection results in higher recruitment of T cells to the tumor site. Both transfected and untransfected cells proliferate at the same rate, both are killed by T cells, and both suppress T cell activity. The model flowchart and equations are shown in Figure 4, and model parameters are listed in Table 1.

**Figure 4.**
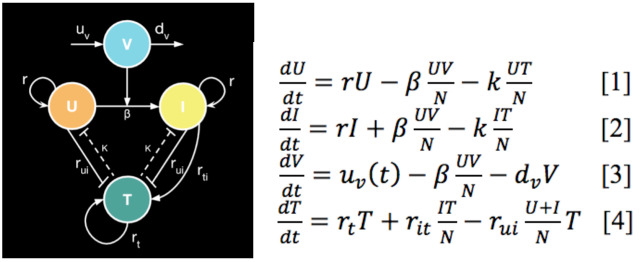
(left) model flowchart; (right) model equations measuring the change in the volume of untransfected cells *dU/dt* [1], transfected cells *dI/dt* [2], the vaccine *dV/dt* [3], and T-cells *dT/dt* [4].

**Table 1.** Parameter values found based on calibration procedures. T-Cell proliferation rate (r_t_) calculated from paper by Hwang et. al. [3].

| Parameter Value |  |
| --- | --- |
| Tumor Cell Net Proliferation | $r = 0.28376$ |
| Transfection Rate | $\beta = 0.534$ |
| Killing Rate | $k = 0.99011$ |
| Decay Rate | $d_v = 0.90015$ |
| T-Cell Net Proliferation | $r_t = 0.46052$ |
| T-Cell Recruitment | $r_{it} = 0.11612$ |
| T-Cell Regulation | $r_{ui} = 1$ |

## VI. DISCUSSION

In this study, various schedules of vaccine injection were tested in order to determine the most effective protocol to reduce tumor burden. We concluded that a single, large, early injection of vaccine yielded the optimal results for tumor reduction. The virtual mice cohort simulations also indicated that this schedule is the most robust in both the objective tumor response, and in the schedule ranking. Therefore, our model showed that this protocol can be applied to many heterogeneous populations.

In certain cases, a single, large injection of the vaccine may not be plausible for laboratory experimentation, thus further studies, both computational and experimental may be needed to test daily smaller injections at the beginning of treatment to mimic the single, large injection that yielded the best results in our simulations.

On the other hand, a single, large injection may be more convenient to use in certain cases. Researchers are hoping to take these treatments to developing countries to treat people in need, and these people would most likely not be able to travel to receive multiple injections of the vaccine. A single, large injection would be necessary to realistically deliver effective treatment.

## ACKNOWLEDGMENT

P.B. and M.K. would like to thank the faculty of HIP IMO for the amazing opportunity to conduct this research.

